# Genetic and Genomic Resources to Study Natural Variation in *Brassica rapa*

**DOI:** 10.1101/2020.06.23.167270

**Authors:** Ping Lou, Scott Woody, Kathleen Greenham, Robert VanBuren, Marivi Colle, Patrick P. Edger, Ryan Sartor, Yakun Zheng, Nathan Levendoski, Jan Lim, Calvin So, Brian Stoveken, Timothy Woody, Jianjun Zhao, Shuxing Shen, Richard M. Amasino, C. Robertson McClung

**Affiliations:** Department of Biological Sciences, Dartmouth College, Hanover, NH, USA; Department of Biochemistry, University of Wisconsin, Madison, WI, USA; Department of Plant and Microbial Biology, University of Minnesota, St. Paul, MN, USA; Department of Horticulture, Michigan State University, East Lansing, MI, 48824, USA; Crop and Soil Sciences, North Carolina State University, Raleigh, NC, 27695, USA; State Key Laboratory of North China Crop Improvement and Regulation, Laboratory of Vegetable Germplasm Innovation and Utilization of Hebei, Collaborative Innovation Center of Vegetable Industry in Hebei, Department of Horticulture, Hebei Agricultural University, Baoding, 071001, China

**Keywords:** *Brassica rapa*, Natural variation, Quantitative Trait Loci, Advanced-Intercross Recombinant Inbred Lines, Seed coat color

## Abstract

The globally important crop *Brassica rapa*, a close relative of Arabidopsis, is an excellent system for modeling our current knowledge of plant growth on a morphologically diverse crop. The long history of *B. rapa* domestication across Asia and Europe provides a unique collection of locally adapted varieties that span large climatic regions with various abiotic and biotic stress tolerance traits. This diverse gene pool provides a rich source of targets with the potential for manipulation towards the enhancement of productivity of crops both within and outside the Brassicaceae. To expand the genetic resources available to study natural variation in *B. rapa*, we constructed an Advanced Intercross Recombinant Inbred (AI-RIL) population using *B. rapa* subsp. *trilocularis* (Yellow Sarson) R500 and the *B. rapa* subsp. *parachinensis* (Cai Xin) variety L58. Our current understanding of genomic structure variation across crops suggests that a single reference genome is insufficient for capturing the genetic diversity within a species. To complement this AI-RIL population and current and future *B. rapa* genomic resources, we generated a *de novo* genome assembly of the *B. rapa* subsp. *trilocularis* (Yellow Sarson) variety R500, the maternal parent of the AI-RIL population. The genetic map for the R500 x L58 population generated using this *de novo* genome was used to map QTL for seed coat color and revealed the improved mapping resolution afforded by this new assembly.

## INTRODUCTION

The globally important crop *Brassica rapa*, a close relative of Arabidopsis, is an excellent system for modeling our current knowledge of plant growth on a morphologically diverse crop. The domestication and spread of *B. rapa* across Europe and Asia provide a diverse collection of varieties locally adapted to widely varying climatic and edaphic regions and subjected to various abiotic and biotic stress challenges. *B. rapa* includes morphologically diverse crops such as turnip, Chinese cabbage, pak choi, leafy vegetables and oilseed (Qi et al., 2017). The demonstrated versatility in *B. rapa* trait cultivation that spans diverse cultural and geographic origins is described in a Chinese almanac (~3000 BCE), ancient Indian texts (~1,500 BCE), and a European link in Babylonia (~722 BCE) (Qi et al., 2017). A whole genome triplication event followed the divergence of *Brassica* from Arabidopsis ~ 23 million years ago (MYA) (Hohmann et al., 2015; Qi et al., 2017). This genome triplication was followed by extensive fractionation (gene loss) (Tang et al., 2012), but likely has contributed to the genetic, morphological, and physiological diversity of *B. rapa* and of the *Brassica* genus in general (Qi et al., 2019). Since the release of the first reference genome in the *B. rapa subsp. pekinensis* Chinese cabbage line Chiifu-401-42 (Wang et al., 2011), *B. rapa* has become an attractive model system because of its complex trait morphology and close relationship with Arabidopsis that facilitates comparative studies.

Since the first *B. rapa* genome release, an updated chromosome-scale assembly (v3.0) with greatly improved contiguity was produced using a combination of long read PacBio data, optical genome maps, and high throughput chromatin conformation capture (Hi-C) (Zhang et al., 2018). A second high-quality, chromosome-scale genome for subsp. *trilocularis* (Yellow Sarson) Z1 was generated using long-read NanoPore data and an optical genome map (Belser et al., 2018). These long-read assemblies have the added advantage of improved detection of transposable elements and mapping of genes located in transposon-rich regions of the genome. The *B. rapa* Z1 assembly identified 20% more Copia elements compared to the reference genome (Belser et al., 2018). Alignment of resequencing data from ~200 *B. rapa* genotypes spanning multiple morphotypes to the Z1 genome supported its utility as a reference for the species (Belser et al., 2018). However, to fully capture the abundant genetic diversity and genomic variation within the species it seems likely that multiple high-quality reference genomes will be needed.

To expand the range of genetic variation available for study, we constructed an Advanced Intercross Recombinant Inbred Line (AI-RIL) population using *B. rapa* subsp. *trilocularis* (Yellow Sarson) R500 as the female and the *B. rapa* subsp. *parachinensis* (Cai Xin) variety L58 as the male parent. The AI-RIL design allows for improved mapping resolution for Quantitative Trait Loci (QTL) identification (Balasubramanian et al., 2009). To facilitate analysis of gene candidates for QTL, we have generated a *de novo* genome assembly for R500. We used this population to map QTL for seed coat color to demonstrate the potential of *de novo* genome assembly to aid the discovery of genes underlying QTL.

## MATERIALS AND METHODS

### R500 Genome assembly and pseudomolecule constructions

Leaf tissue from the first fully developed leaves of 3-week old R500 plants was harvested following 24 h of dark treatment. Tissue was flash frozen in liquid nitrogen and stored at −80°C. A total of ~90 g of tissue was shipped to the Arizona Genomics Institute at the University of Arizona for high molecular weight genomic DNA extraction and PacBio sequencing.

R500 PacBio reads (NCBI Sequence Read Archive [SRA] SRR12035043) were error-corrected and assembled using Falcon (v0.2.2) (Chin et al., 2016). Parameters for Falcon were modified as follows in the configuration file: pa_HPCdaligner_option = -v -dal128 -t16 -e.70 -l1000 -s1500 ovlp_HPCdaligner_option = -v -dal128 -t32 -h60 -e.96 -l500 -s1500. falcon_sense_option = -output-multi --min-idt 0.75 --min-cov 6 --max-n-read 250. The resulting graph-based assembly was visualized in Bandage (Wick et al., 2015) to verify assembly quality. Draft Falcon-based contigs were polished to remove residual errors with Pilon (v1.22) using 69.8x coverage of Illumina 150 bp paired-end libraries prepared from R500 gDNA. Raw Illumina reads (NCBI SRA - SRR496614) were quality-filtered using Trimmomatic (Bolger et al., 2014) with default parameters and aligned to the Falcon based contigs using bowtie2 v2.3.0 (Langmead and Salzberg, 2012) with default parameters. The total alignment rate of the Illumina data was 96.3%, suggesting our assembly was nearly complete. The following Pilon parameters were modified: --flank 7, --K 49, and --mindepth 10 and all other parameters were left as default. Pilon was run reiteratively four times with realignment to the updated assembly for each pass. The fourth pass corrected few additional InDel and SNP based errors suggesting the assembly was sufficiently polished.

The Pilon based contigs were anchored into a chromosome scale assembly using a high-density genetic map that we constructed from R500/IMB211 SNPs identified through RNA-seq analysis of individual RIL from an R500 x IMB211 RIL population (Markelz et al., 2017) (Supplemental Table S1). Contigs were anchored to the pseudomolecules if they contained a minimum of three markers and contigs were ordered based on marker orientation in the genetic map. Contigs were stitched together with addition of interstitial 10,000 N spacer sequences.

### R500 Genome Annotation

The *B. rapa* R500 genome was annotated using the MAKER annotation pipeline (Campbell et al., 2014). Transcript and protein evidence used in the annotation included protein sequences downloaded from Araport11 and Phytozome12 plant databases, *B. rapa* expressed sequence tags (EST) from NCBI, and transcriptome data downloaded from NCBI and generated from different *B. rapa* leaf tissues under drought treatments and assembled with StringTie (Pertea et al., 2015) or Trinity (Haas et al., 2013). Repetitive regions in the genome were masked using a custom repeat library and Repbase (Jurka et al., 2005) through Repeatmasker (Smit et al., 1996). *Ab initio* gene prediction was performed using the gene predictors SNAP (Korf, 2004) and Augustus (Stanke and Waack, 2003). The resulting MAKER gene set was filtered to select gene models containing Pfam domain and annotation edit distance (AED) < 1.0 and scanned for transposase coding regions. The amino acid sequence of predicted genes was searched (BLASTP, 1e-10) against a transposase database (Campbell et al., 2014). The alignment between the genes and the transposases was further filtered for those caused by the presence of sequences with low complexity. The total length of genes matching transposases was calculated based on the output from the search. If more than 30% of gene length aligned to the transposases, the gene was removed from the gene set. Furthermore, to assess the completeness of annotation, the *B. rapa* Maker gene set was searched against the Benchmarking Universal Single-Copy Orthologs (BUSCO v.2) (Simão et al., 2015) plant dataset (embryophyta_odb9). We identified a total of 42,381 protein coding genes. To identify Arabidopsis orthologs, we first ran a BLAST search using protein coding sequences and pulled out the top 3 hits using an e-value cutoff of 0.001. To define the Arabidopsis ortholog, the top BLAST hit was first selected; if the ortholog was not located in the correct syntenic block (Parkin et al., 2005; Schranz et al., 2006; Zhang et al., 2018), we screened all candidate orthologs based on gene structure and syntenic block. This resulted in 35,157 *B. rapa* genes with predicted Arabidopsis orthologs (Supplemental Table S2). Based on chromosomal positioning we have matched the R500V1.1 gene annotations with the Chiifu v1 gene annotations used in the NCBI and *EnsemblPlants* databases (Supplemental Table S3).

Long terminal repeat (LTR) retrotransposons in the *B. rapa* R500 genome were identified using LTRharvest (Ellinghaus et al., 2008) and LTR_finder (Xu and Wang, 2007). A non-redundant LTR library was produced by LTR_retriever (Ou and Jiang, 2018). Miniature inverted transposable elements (MITEs) were identified using MITE-Hunter (Han and Wessler, 2010) manually checked for target site duplications (TSD) and terminal inverted repeats (TIR) and classified into superfamilies. Those with ambiguous TSD and TIR were classified as “unknowns.” Using the MITE and LTR libraries, the *B. rapa* genome was masked using Repeatmasker (Smit and Hubley, 2008). The masked genome was further mined for repetitive elements using Repeatmodeler (Smit and Hubley, 2008). The LTR libraries and corresponding location in the genome is provided as Supplemental File S1. The repeats were then categorized into two groups based on whether they had homology to classified families. Those without identities were searched against the transposase database and if they had a match, they were considered a transposon. The repeats were then filtered to exclude gene fragments using ProtExcluder (Campbell et al., 2014) and summarized using the “fam_coverage.pl” script in the LTR_retriever package (Ou and Jiang, 2018).

We supplemented the InterPro-based annotations (Mitchell et al., 2019) with the Arabidopsis homolog annotations resulting in roughly 38,000 annotated genes. For InterPro, the GO evidence codes were not included since all of them are IEA (inferred from electronic annotation). The “InterPro_ID” listed in Supplemental Table S4 is the InterPro protein domain that was used for the annotation. For genes that were annotated based on Arabidopsis homology, the Arabidopsis ortholog was used to infer the term. For the GOslim annotations, we used the GO consortium “owltools” (https://github.com/owlcollab/owltools/wiki) software “map2slim” program to map the GO annotations to the Plant GOslim ontology (Supplemental Table S5). Finally, to associate KEGG terms, we used KAAS (KEGG automated annotation server) that outputs the KEGG ontology number (Moriya et al., 2007). This was then matched to the Pathways and Enzymes (Supplemental Table S6).

### Construction of R500 x L58 population

We constructed an Advanced Intercross-Recombinant Inbred Line (AI-RIL) population using *B. rapa* subsp. *trilocularis* (Yellow Sarson) R500 as the female and the *B. rapa* subsp. *parachinensis* (Cai Xin) variety L58 (Zhao et al., 2010) as the male parent. The breeding program was broadly analogous to that described for the creation of AI-RILs in Arabidopsis (Balasubramanian et al., 2009). 18 F2 lineages were used to initiate three successive generations of pairwise, nonredundant intercrosses (IC1F1 – IC3F1 generations in our nomenclature). Two progeny seeds of each IC*n*F1 generation were planted and grown in successive rounds of intercrosses to maintain the IC breeding populations at 36 individuals. IC3F1 plants were self-pollinated and 196 IC3F2 lines were used to establish independent lineages that were advanced through 6 generations of selfing with single seed descent (*s*2 – *s*8) followed by bulking (Figure 1). Freshly harvested seeds were placed on a bed of moist soil (MetroMix 360), covered with a thin layer of coarse grade vermiculite, and watered to moisten the vermiculite top-coat. Neither stratification nor vernalization is required. Plants were grown in a greenhouse under 16 h light at 23°C and 8 h dark at 20°C. Supplemental light was provided as needed by use of high-pressure sodium lamps.

**Figure 1.**
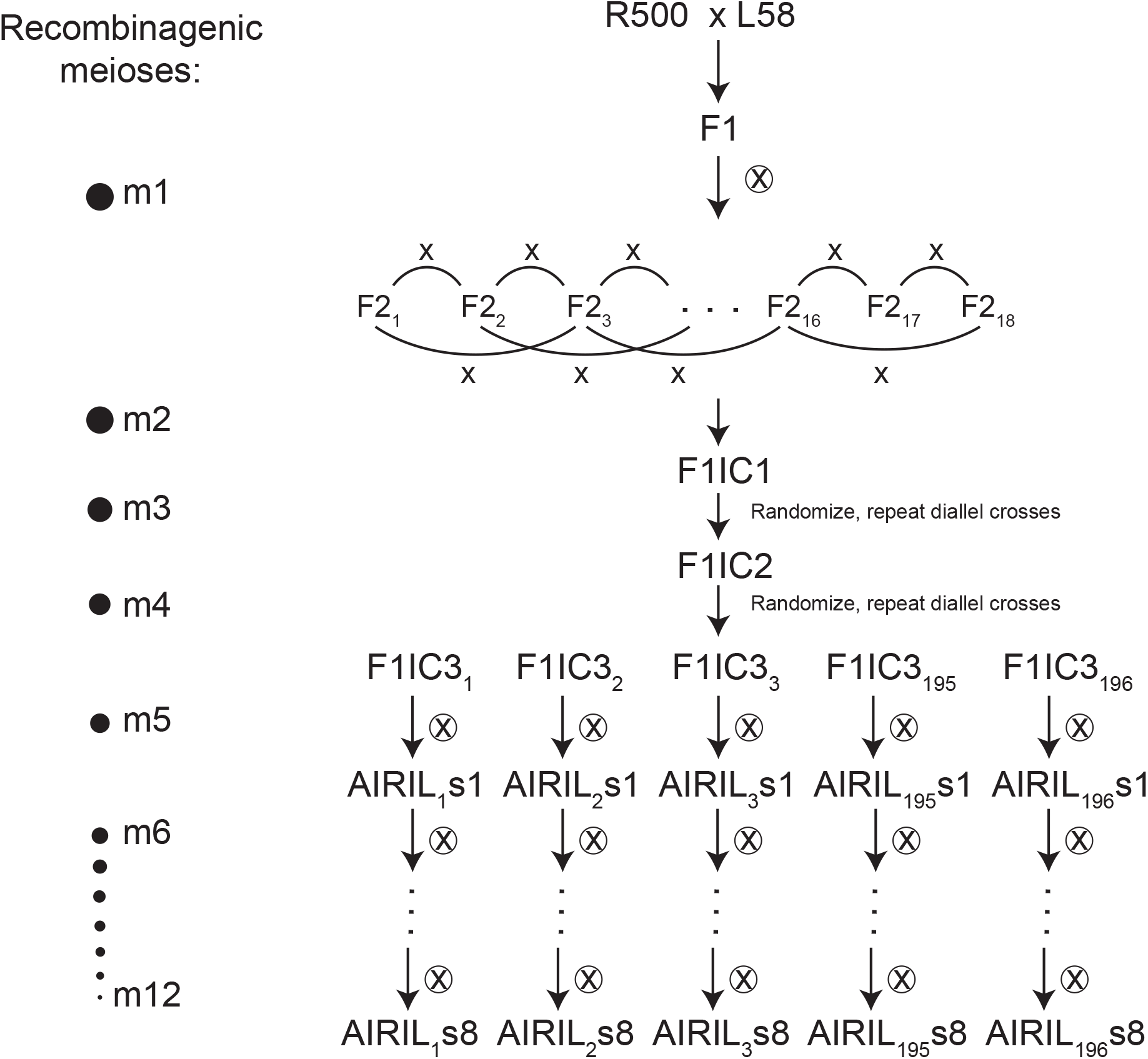
Development of a R500 x L58 Advanced Intercross Recombinant Inbred Line (AI-RIL) population for quantitative genetics. The population was founded by crosses between the self-compatible historically inbred R500 variety as the maternal parent and L58 as the male parent using the crossing strategy shown. Intercross generations (IC) were founded from 18 F2 segregants. Random 2x pairwise crosses were advanced to IC3F2 followed by 8 selfing generations. 196 discrete AI-RIL were propagated by single seed descent. Pooled tissue from young leaves of s7 plants was used for GBS.

### Genotype-by-Sequencing (GBS) of the R500 x L58 population

Pooled young leaves of three individual plants from each of 186 lines of the S7 generation were sent to the University of Minnesota Genomics Center for DNA extraction and GBS. DNA was digested with *ApeK1* followed by Illumina adapter and barcode ligation. Libraries were sequenced on one lane of a NovaSeq 1X100 SP Flowcell for ~2M reads/sample. Sequencing data have been deposited in the NCBI SRA under PRJNA625700.

### Constructing a high-density genetic map for R500 x L58

Raw reads were de-multiplexed, trimmed to 70 bases and filtered with a quality score cutoff of 28 using FASTX-Toolkit (v0.0.13.2; http://hannonlab.cshl.edu/fastx_toolkit/index.html). Filtered reads from R500 x L58 were mapped to the R500 genome v1.2 (https://genomevolution.org/coge/) using BWA (Li and Durbin, 2009). SNPs were called using GATK v2.8 with the parameters: -T UnifiedGenotyper --genotyping-mode DISCOVERY (McKenna et al., 2010). SNPs from GATK were filtered using the VCFtools software v0.1.12a (Danecek et al., 2011) with the following parameters: --remove-indels --maf 0.05 --mac 10 -- min-alleles 2 --max-alleles 2 --max-missing-count 30. All heterozygous SNPs were treated as missing. SNPbinner (Gonda et al., 2019) was used for identifying the crossover events (crosspoints) and to generate a high-resolution bin-based genetic map of the RIL population; after selecting tagging SNPs from each bin, 1109 SNPs from 184 lines were selected to construct the draft linkage map. The linkage map was generated using onemap v1.0-1 (Margarido et al., 2007) in R (Team, 2018) with manually imputed marker data based on a hidden markov model in biallelic population (Lincoln and Lander, 1992) and then corrected for genotyping errors using R/QTL, Calc.errorlod function. The final round of map construction was performed again using onemap and linkage groups were assigned to chromosomes based on the R500 reference genome.

### Seed Coat Color

Approximately 100 seeds from each of the two parental lines and 184 AI-RILs were used to evaluate seed coat color. Seeds from each line were placed in a white plastic weigh boat and photographed with a Canon EOS 450D camera with fixed lens and shutter speed under controlled light conditions. Images were imported into MATLAB and average RGB values were obtained for identically-sized Regions of Interest (ROI) to yield a quantitative representation of seed coat color for QTL analysis. Lines with limited seed number or with non-uniform seed coat color were recorded as missing values.

### QTL analysis

QTL analysis was conducted by using R/QTL package 1.41-6 (Broman et al., 2003) in an RStudio environment running R version 3.4.1 (https://cran.r-project.org/bin/windows/base/). A full transcript of our analyses suitable for use by those who might be interested to repeat or to refine our analysis is provided in Supplemental File S2. Briefly, we used phenotypic and genotypic data provided in Supplemental Table S7 (R500 x L58 AI-RIL population) as input to R/QTL, followed by invocation of jittermap with parameter amount=1e-6, convert2riself, and calc genoprob (step=0.5, error.prob=0.001 functions. The scanone function (method=“em”) was used to identify primary QTL under a single QTL model, followed by composite interval mapping (method=“cim”) using SNP close to the QTL peak as cofactors. LOD thresholds (*p* < 0.05)_were determined through 1000 data permutations, and the map position and extent of statistically significant interval was determined by using lodint at 1.5.

## RESULTS

### R500 genome assembly

A high-quality reference genome of the *B. rapa subsp. trilocularis* (Yellow Sarson) R500 was generated using a PacBio based, single-molecule, real-time (SMRT) sequencing approach. In total, we sequenced 28 SMRT cells (2 at 4h, 26 at 6h) with a subread N50 of 16.6 kb and generated 2.0 million raw PacBio reads collectively spanning 29.5 Gb to achieve 54.6x coverage of the 540 Mb genome. Falcon (v0.2.2) was used for the assembly and polished with Pilon (v1.22) using 69.8x coverage of Illumina 150 bp paired-end DNA-sequencing data from R500. The Pilon based contigs were anchored into chromosomes using an updated high-density genetic map that was constructed from GBS-based variants of an R500 x IMB211 population (Supplemental Table S1) (Markelz et al., 2017). The R500 genome assembly V1.2 is available on CoGe (https://genomevolution.org/coge/). Assembly statistics are reported in Table 1. In total, 127 contigs collectively spanning 280.5 Mb (or 78.8% of the assembly) were anchored and oriented into ten chromosomes. The vast majority (42381 of 45538; ~93%) of gene models were anchored to chromosomes. Gene density was lowest near the centromeres and higher along the chromosomal arms (Figure 2). The Falcon based assembly has 1,753 contigs spanning 356 Mb with a contig N50 of 3.9 Mb and N90 of 90 kb. The longest contig is 24.5 Mb and spans a full arm of chromosome A03. The only complex region in the assembly graph spans several high copy number long terminal repeat retrotransposons and rRNA repeats in the nucleolus organizer region. LTR transposable elements were found throughout the chromosomes (Figure 2).

**Figure 2.**
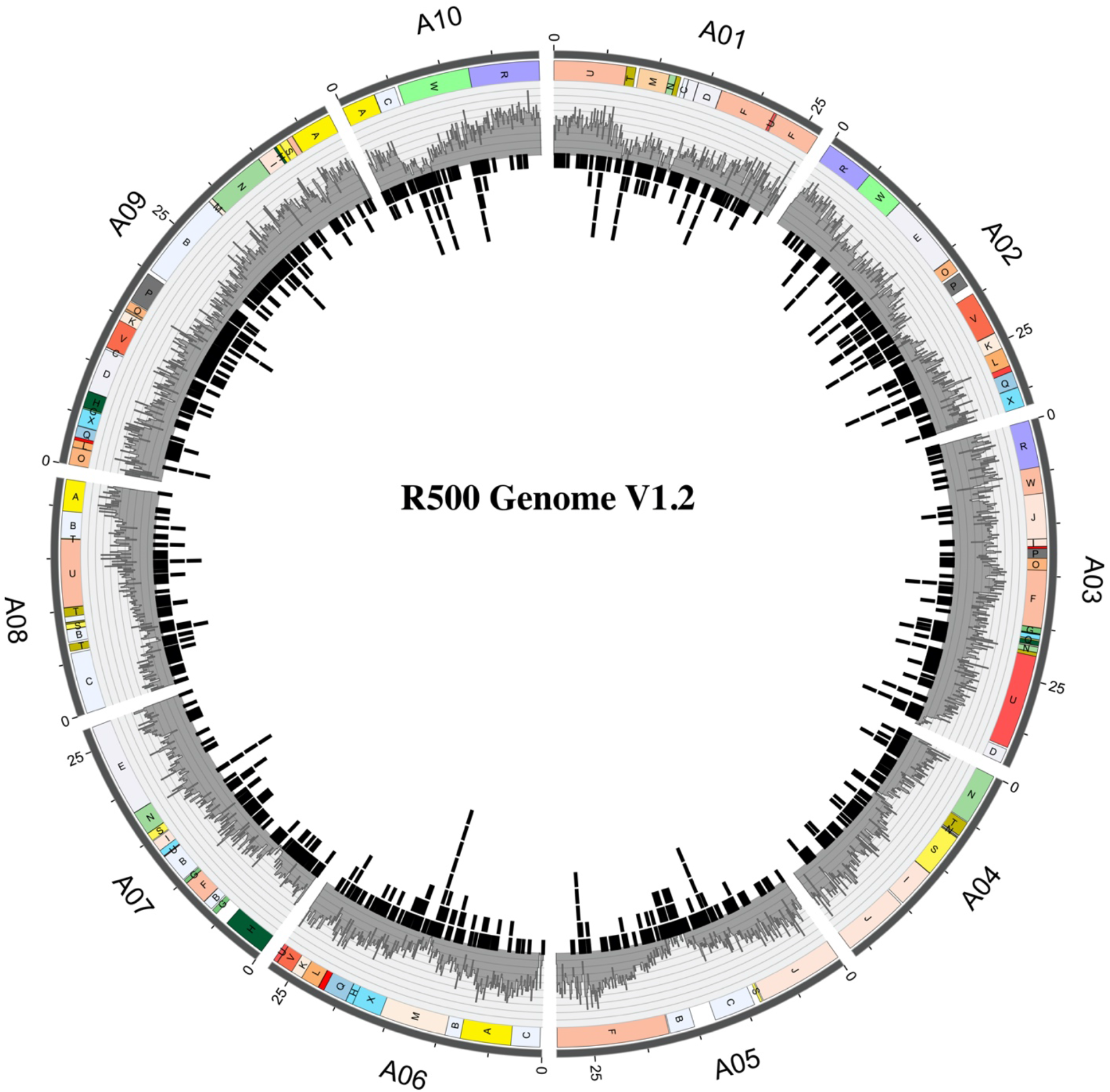
Circos plot illustrating the *B. rapa* R500 genome V1.2. The ten chromosomes are displayed in the outer circle, with chromosomal length in Mbp shown in 5 Mbp increments). Blocks syntenic with Arabidopsis (Parkin et al., 2005; Schranz et al., 2006; Zhang et al., 2018) are indicated by the colored boxes labelled A through X in the circle immediately inside the chromosomes. The third circle from the outside provides in gray gene distribution as a density histogram (numbers of genes per 100 Kbp, ranging from 0-40/100Kbp). The innermost circle shows LTR distribution.

**Table 1:** *Brassica rapa* subsp. *trilocularis* (Yellow Sarson) R500V1.2 genome assembly statistics.

|  |  |
| --- | --- |
| Number of contigs | 1753 |
| Longest contig | 24.5 Mb |
| Contig N50 | 3.9 Mb |
| Contig N90 | 90 Kb |
| Unanchored | 75 MB |
| Total Size | 356 Mb |
| % Anchored | 78.80 |

Comparison of the *B. rapa* R500 V1.2 and Chiifu V3.0 (Zhang et al., 2018) pseudomolecules revealed 1:1 collinearity across all 10 chromosome pairs with several notable large-scale inversions and structural differences (Figure 3A). Differences may be due to errors in genome assembly and anchoring or they could represent true structural variation between these diverse accessions. We note that the discontinuities typically fall either at chromosome ends or internally, associated with centromeres, and both regions are associated with repetitive DNA, which can be challenging to assemble. Similarly, the *B. rapa* R500 and Yellow Sarsen Z1 (Belser et al., 2018) pseudomolecules revealed overall collinearity with several differences, most of which are associated with regions in the Z1 assembly not represented in the R500 assembly (Figure 3B). We speculate that these are probably attributable to the advanced assembly techniques, such as nanopore long read sequencing and optical mapping, allowing incorporation of a greater proportion of repetitive sequences. The remaining unmapped R500 contigs likely correspond to highly repetitive pericentromeric or telomeric regions which have low marker density and low recombination rates, hindering their accurate anchoring.

**Figure 3.**
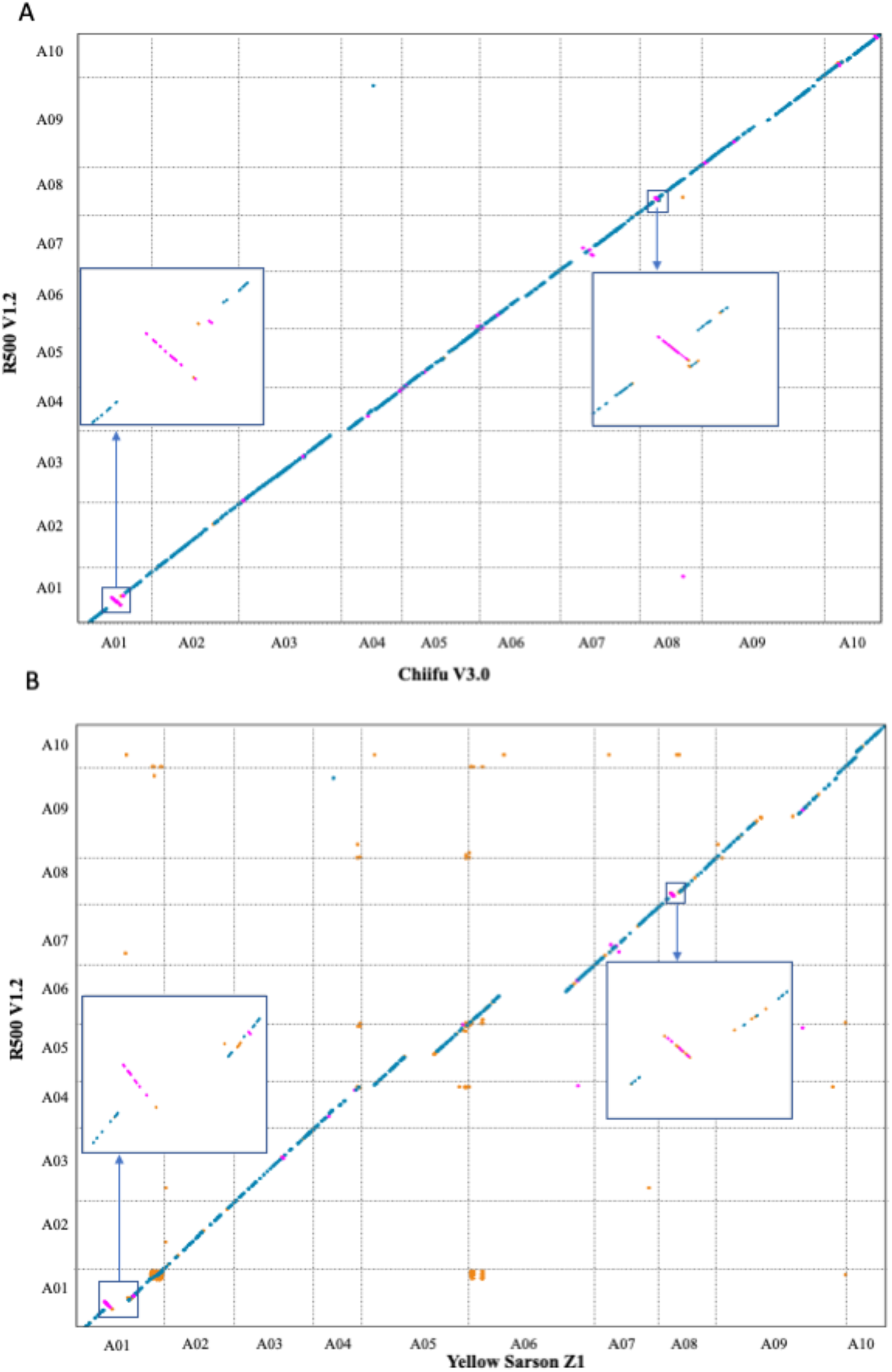
Macrosyntenic comparisons of the R500 V1.2 assembly with Chiifu V3.0 (A) and Yellow sarson Z1 (B). Each black dot represents a syntenic gene pair between two genomes and deviations from diagonal lines between chromosome pairs denote structural variations, inversions, or assembly errors between the genomes. Examples of these deviations are shown in the highlighted boxes.

### Genetic map of the R500 x L58 AI-RIL population

The breeding program used to construct the AI-RIL populations is broadly analogous to that described for the creation of AI-RILs in Arabidopsis (Balasubramanian et al., 2009). To generate high density genetic linkage maps for R500 x L58 population, Genotype-by-Sequencing (GBS) was performed on the s7 generation. The genetic map is provided in Figure 4 and Supplemental Table S7. One advantage of the advanced intercrossing strategy employed is increased resolution due to increased recombination density (Balasubramanian et al., 2006). Consistent with this expectation, we determined the number of crossovers per line to be 42.91 ± 14.64, more than double that (19.79 ± 4.61) of a second *B. rapa* RIL population (Iniguez-Luy et al., 2009) generated without the intercrossing steps. The advanced intercrossing design led to expansion of the genetic map, which contains recombination events corresponding to 481 kb/cM. Both parents, R500 and L58, and 186 AI-RIL lines used in genetic map construction have been deposited with the Arabidopsis Biological Resource Center (Ohio State University; https://abrc.osu.edu) as accession numbers CS28987, CS28988, and CS99437 - CS99622, respectively.

**Figure 4.**
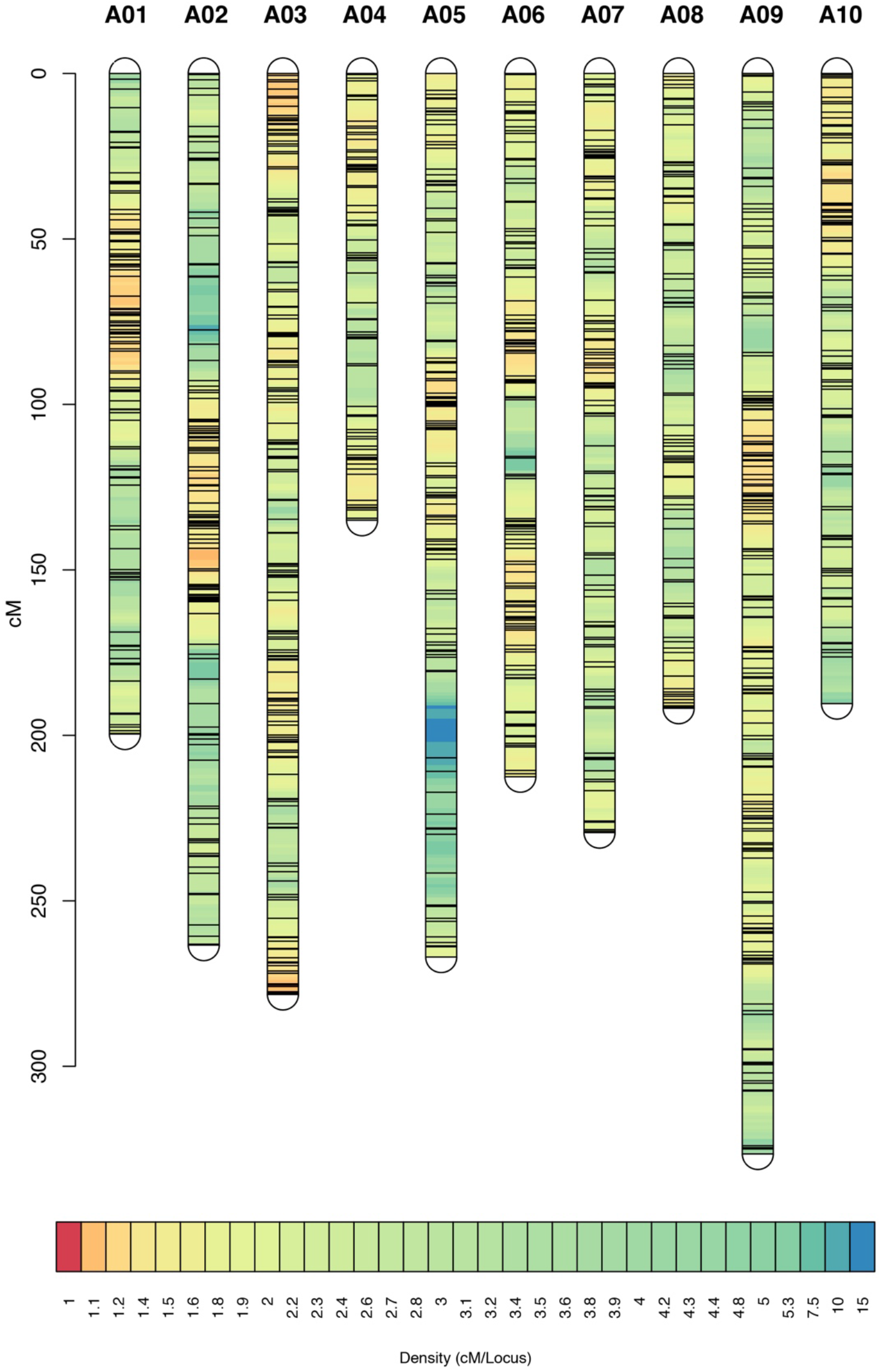
Density plot of SNP markers in the R500 x L58 AI-RIL population. Marker locations for all 10 *B. rapa* chromosomes represented as a Density plot, modified from LinkageMapView (Ouellette et al., 2017). The scale on the left represents the map position in centiMorgans (cM).

### Natural Variation in Seed Coat Color (SCC)

We sought to validate the utility of the new RIL population by testing for natural variation in seed coat color. R500, the maternal parent, has yellow seeds whereas L58 has dark brown seeds. We detected a strong QTL on chromosome A09, accounting for ~37% of the variance in seed coat colors in the R500 x L58 population. (Figure 5A, Table 2, Supplemental Table S7). *TRANSPARENT TESTA8* (*TT8*), which encodes a bHLH transcription factor that positively regulates proanthocyanin biosynthetic pathways (Nesi et al., 2000), colocalizes with the QTL interval and constitutes a strong candidate for this seed coat color QTL. Although *B. rapa* has undergone a whole genome triplication since its separation from Arabidopsis, there has been considerable gene loss, termed fractionation, since the triplication (Town et al., 2006; Wang et al., 2011) and *TT8* is present only in a single copy in *B. rapa*. Accordingly, we interrogated our new genome assembly of R500 as well as the resequencing data for L58 (Zhang et al., 2020) and found a helitron transposable element inserted in the R500 *TT8* locus whereas the L58 *TT8* allele was intact and predicted to encode functional protein (Figure 5B). Thus, we conclude that *TT8* is a strong candidate locus responsible for this seed coat color QTL. This highlights the importance of having genome assemblies of parental lines to improve gene discovery of identified QTLs.

**Figure 5.**
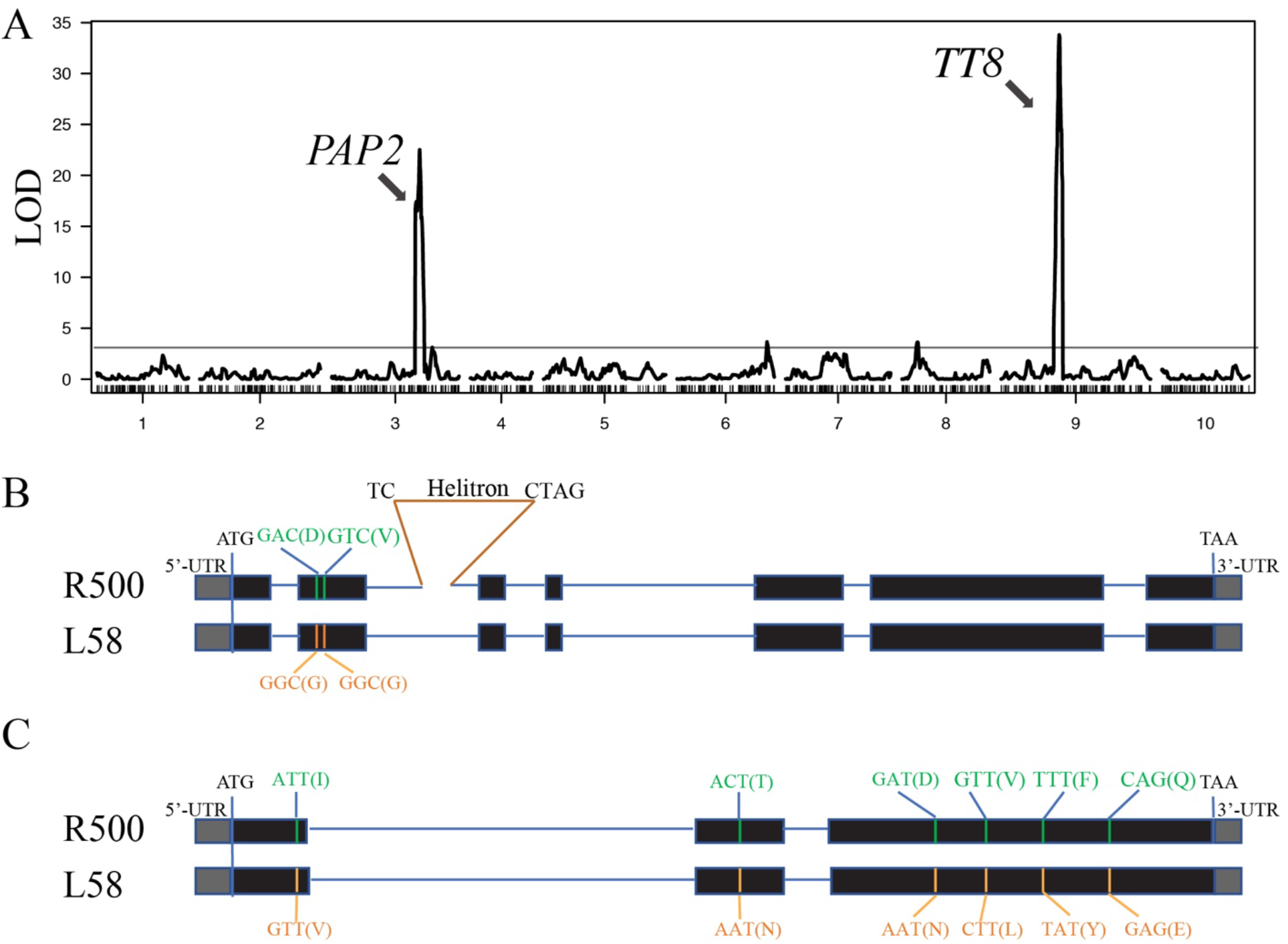
QTL mapping results for seed coat color in the R500 x L58 AI-RIL population. A. The R/qtl program was used to identify a major QTL on chromosome A09 for seed coat color that accounts for ~37% of the phenotypic variation in the population. Underlying this QTL is the *TT8* locus that has been shown to be responsible for seed coat color in another yellow-seeded *B. rapa* (Li et al., 2012). A second major QTL for seed coat color that accounts for ~32% of the phenotypic variation in the population was identified on chromosome A03. Underlying this QTL is the candidate gene *PAP2*. Horizontal line indicates significance threshold. B. Cartoon of the R500 and L58 *TT8* alleles, with exons indicated by boxes (5’ and 3’ UTRs are filled with gray and coding sequences filled with black) and introns indicated by horizontal lines. Two single nucleotide polymorphisms resulting in amino acid substitutions are indicated, as is the site of insertion of a Helitron transposable element into the second intron of the R500 *TT8* sequence. C. Cartoon of the R500 and L58 *PAP2(A03)* alleles, with exons indicated by boxes (5’ and 3’ UTRs are filled with gray and coding sequences filled with black) and introns indicated by horizontal lines. Single nucleotide polymorphisms resulting in amino acid substitutions are indicated, although it is not known whether any of these substitutions affect PAP2 activity.

**Table 2.** *SEED COAT COLOR (SCC)* Quantitative Trait Loci in a Yellow Sarson (R500) x Cai Xin (L58) Advanced Intercross-Recombinant Inbred Line (AI-RIL) population.

| Chromosome | Position<br>(cM) | LOD | Variance<br>(%) | Cofactors | Confidence Interval | Candidate<br>genes |
| --- | --- | --- | --- | --- | --- | --- |
| A09 | 125 | 43.46 | 36.79 | A09_17.56 | A09_17.14-A09_21.43 | *TT8* |
| A03 | 195 | 22.74 | 31.77 | A03_20.08 | A03_17.8-A03_20.28 | *PAP2* |

We also detected a QTL on chromosome A03 that accounted for ~32% of the variance in seed coat color (Table 2, Figure 5A, Supplemental Table S7). Again, a strong candidate, *PRODUCTION OF ANTHOCYANIN PIGMENT2* (*PAP2*(*A03*)) mapped to this chromosomal region. In Arabidopsis, *PAP1* and to a lesser extent its close homologue *PAP2* encode R2R3 Myb domain transcription factors (AtMyb75 and AtMyb90, respectively) that have been shown to be important for light dependent accumulation of anthocyanin (Cominelli et al., 2008). *B. rapa* has three genes homologous to the Arabidopsis *PAP* genes. By simple sequence similarity it is not possible to determine unequivocally whether they are true orthologues to *AtPAP1* or *AtPAP2*. However, by synteny (Parkin et al., 2005; Schranz et al., 2006; Zhang et al., 2018), it is clear that two of the *B. rapa* loci, on Chr. A02 and A07, are orthologous to *AtPAP2* (Supplemental Figure S1A). The third *B. rapa PAP* locus is not in a region of Chr. A03 syntenic with either *AtPAP1* or *AtPAP2* but is slightly greater in amino acid identity to *At PAP2* than to *AtPAP1* (Supplemental Table S8A). This argues that *B. rapa PAP*(*A03*) represents a *PAP2* ortholog, which we call *PAP2*(*A03*).

Accordingly, we will refer to each of these *B. rapa* genes as *PAP2(A02), PAP2(A03)*, and *PAP2(A07)*. The R500 and L58 alleles of *PAP2(A03)* can be distinguished by a number of SNPs, including several (6) predicted to result in changes to the amino acid sequence (Figure 5C, Supplemental Figures S1B and S1C, Supplemental Table S8B). At four of these amino acid positions (amino acid positions 36, 150, 168, and 210) the L58 residue is conserved with Arabidopsis PAP1 and PAP2 as well as with *B. rapa* PAP2(A02) and PAP2(A07) whereas the R500 allele has a different residue (V36I, N150D, L168V, and E210Q, where the first residue is that of L58 and the other PAP proteins and the second residue is that of R500 PAP2(A03)). This is consistent with the hypothesis that one or more of these substitutions in the R500 PAP3(A03) result in loss of function, which would be consistent with the lack of seed coat pigmentation in the yellow R500 seeds. In particular, we note that residue 36 is in the R2-Myb domain, which is required for DNA-binding activity. However, at this time we have no functional data supporting that these amino acid substitutions have a major impact on the function of the R500 PAP2(A03) protein. Thus, although *PAP2(A03)* is a strong candidate for this QTL, confirmation will require additional experimentation.

## DISCUSSION

### R500 *de novo* genome assembly

To improve the mapping resolution of identified QTL and facilitate follow-up studies we have generated a *de novo* genome assembly for R500. The choice of reference genome significantly influences gene expression and isoform identification even among closely related species (Slabaugh et al., 2019). As a result, identifying causal loci for QTL can be extremely challenging when genetic maps are created based on a reference genome that does not reflect the varieties represented in the population. Access to the R500 genome assembly and the L58 sequence data (Zhang et al., 2020) improved the quality of the genetic maps and facilitated the identification of candidate genes for mapped QTL.

### Natural variation in seed coat color (SCC)

To demonstrate the utility of the R500 x L58 AI-RIL population for exploring natural variation, we chose to map QTL for one previously characterized trait. Several groups have identified QTL for seed coat color in these genomic regions of *B. rapa* (Kebede et al., 2012; Li et al., 2012; Rahman et al., 2014; Wang et al., 2016; Zhao et al., 2019) and *B. juncea* (Padmaja et al., 2014). Colocalized with the A09 SCC-QTL was *TT8*, a previously identified bHLH transcription factor that regulates proanthocyanin biosynthesis (Nesi et al., 2000). Using our R500 assembly, we found a helitron transposable element insertion in the R500 allele of *TT8* strongly suggesting inactivation whereas L58 showed intact alleles for *TT8*. These data are consistent with *TT8* being responsible for the SCC QTL on chromosome A09 in these populations. Our discovery of a transposable element disrupting the *TT8* locus in R500 is consistent with an earlier study (Li et al., 2012) that also mapped a major SCC-QTL peak on chromosome A09 among RILs derived from a cross of 3H219 (black-seeded parent) as a donor to Yellow sarson (yellow-seeded parent). They further showed that *TT8* was inactivated by the insertion of a helitron transposable element whereas that *TT8* allele of the black-seeded parent was intact and fully functional. The genetic relationship between their Yellow sarson variety and R500 is not known.

We identified a second SCC QTL on Chromosome A03 for which *B. rapa PAP2* is a strong candidate. *PAP1* and *PAP2* encode Myb-domain transcription factors important for the expression of flavonoid biosynthetic genes. Overexpression of either *PAP1* or *PAP2* greatly enhances anthocyanin pigmentation in Arabidopsis, tobacco (Borevitz et al., 2000), and tomato (Li et al., 2018). Although the R500 and L58 alleles of *PAP2(A03)* can be distinguished by a number of SNPs, in the absence of functional data we cannot confirm the hypothesis that *PAP2(A03)* is responsible for this QTL.

In summary, we provide a new whole-genome assembly as well as a new AI-RIL population suitable for QTL analysis of natural variation between two distinct *B. rapa* morphotypes. These resources should facilitate efforts to understand the genetic bases of the morphological, physiological, and biochemical differences among these diverse varieties.

## Acknowledgements

We thank Guusje Bonnema for the *Brassica rapa* doubled haploid line L58. This work was supported by National **Science** Foundation grants IOS-1202779 to K.G., IOS-1711662 to R.S., and IOS-1547796 the R.M.A. and C.R.M., and by the Rural Development Administration, Republic of Korea Next Generation BioGreen 21 grant number SSAC PJ01327306 to C.R.M.

## Author contributions

P.L., S.W., K.G., R.V., P.P.E., J.Z., R.M.A., and C.R.M. designed the research; P.L., S.W., K.G., R.V., P.P.E., M.C., R.S., Y.Z., N.L., J.L., C.S., B.S., and T.W. performed research; P.L., S.W., K.G., R.V., P.P.E., M.C., and R.S. analyzed data; P.L., S.W., K.G., R.V., P.P.E., R.S., J.Z., R.M.A., and C.R.M wrote the paper.

## Supplemental Materials

**Supplemental Figure S1. (A)** Microsynteny analysis of members of the *PAP* Gene Family in Arabidopsis and *Brassica rapa. B. rapa* genes retained after fractionation in each of the genomic regions syntenic to the Arabidopsis PAP genes are indicated. **(B)** Multiple alignment of PAP proteins of *Arabidopsis* and *Brassica rapa*. Amino acid differences between the R500 and L58 PAP(A03) variants are colored, with highly conserved residues in red and dissimilar residues in yellow or cyan. *, :, and . indicates fully, strongly, and weakly conserved, respectively. **(C)** Nucleotide sequence alignment of R500 (upper) and L58 (lower) alleles of the *PAP2(A03)* (BraA03g40160R) gene. Exonic sequences are highlighted in cyan and intronic sequences are not highlighted. Nucleotide polymorphisms are indicated by red, with substitutions indicated by # and indels indicated by filled boxes. Asterisks mark 10 bp intervals.

**Supplemental Table S1.** (A) Genetic map positions of Single Nucleotide Polymorphism (SNP) Markers and (B) marker inheritance in 121 lines from the R500 x IMB211 Recombinant Inbred Line population (Markelz et al., 2017) used to construct the high-density genetic map onto which the Pilon based contigs were anchored into a chromosome scale assembly of the *B. rapa* R500 genome. (B) Single Nucleotide Polymorphism (SNP) marker inheritance in 121 lines from the R500 x IMB211 Recombinant Inbred Line population (Markelz et al., 2017) used to construct the high-density genetic map onto which the Pilon based contigs were anchored into a chromosome scale assembly of the *B. rapa* R500 genome.

**Supplemental Table S2.** List of predicted protein-coding genes in *B. rapa* R500 V 1.2. Start and end positions for each predicted coding sequence are indicated. Strand indicates the mRNA-like strand of the DNA. Block refers to the syntenic block in the *A. thaliana* genome (Parkin et al., 2005; Zhang et al., 2018).

**Supplemental Table S3.** Correspondence of the R500 V1.2 predicted protein-coding gene annotations with the Chiifu V1 gene annotations used in the NCBI and *EnsemblPlants* databases.

**Supplemental Table S4.** Gene Ontology (GO) terms for genes in the *B. rapa* R500 genome. One or more GO terms were assigned to 38,197 unique gene annotations. The “Gene_ID” field lists *B. rapa* R500 gene accessions. The “GO_ID” field lists the GO term (category) associated with each gene. The “InterPro_ID” field lists accession numbers for functionally characterized InterPro protein domains (see https://www.ebi.ac.uk/interpro). InterPro domains are used to assign GO terms to uncharacterized genes. If no InterPro domain is found, this field has “NA” and the closest Arabidopsis homolog is used to assign GO terms. In this case, the field “At_Homolog” lists the closest Arabidopsis protein match for each gene. The field “Assignment_Type” has an entry of either “InterProScan” or “AtHomology” and specifies which method was used to assign each GO term. The field “GO_Term_Name” lists the short name for each term. The field “GO_Ontology” specifies which of the 3 ontologies (Molecular Function, Biological Process or Cellular Component) each term comes from. The field “GO_Term_Definition” provides a longer description for each term.

**Supplemental Table S5.** Gene Ontology (GO) Slim terms for genes in the R500 genome. GO Slim is a greatly reduced set of higher level (more broad) Gene Ontology terms. One or more of only 97 GO Slim terms is assigned to 38,199 unique gene annotations. The “Gene_ID” field list R500 gene accessions. The “GO_ID(GOSlim)” field lists the GO Slim term (category) associated with each gene. The field “GO_Term_Name” lists the short name for each term. The field “GO_Ontology” specifies which of the 3 ontologies (Molecular Function, Biological Process or Cellular Component) each term comes from. The field “GO_Term_Definition” provides a longer description for each term.

**Supplemental Table S6.** Kyoto Encyclopedia of Genes and Genomes (KEGG) orthology-based functional annotations for genes in the *B. rapa* R500 genome. KEGG uses experimentally confirmed protein functions across various organisms to assign function to unknown proteins based on orthology (see https://www.genome.jp/kegg/ for details and descriptions). Each of 12,428 genes are annotated with one of 3,516 KEGG orthology terms. The field “Gene_Name” lists R500 gene accessions. The field “KEGG_ONTOLOGY_NUMBER” lists the KEGG Orthology term associated with each gene. This is an accession for a functional annotation category. The field “KEGG_PATHWAY”, when available, lists one or more KEGG Pathway accessions separated by “|”. A KEGG pathway is a defined biochemical pathway in which the gene may be involved. The field “KEGG_ENZYME”, when available, lists one or more KEGG enzymes. This is a list of potential known enzymes that each gene may encode.

**Supplemental Table S7. (A)** Genetic map positions of Single Nucleotide Polymorphism (SNP) Markers for 186 lines from the R500 x L58 Advanced Intercross-Recombinant Inbred Line population was used to construct the high-density genetic map shown in **Figure 2**. Seed coat color, measured as average RGB values for an image of 100 seeds, was used to map the QTL as summarized in **Figure 4A** and **Table 2**. Parental values are included last. (B) Genomic positions of Single Nucleotide Polymorphism (SNP) markers selected for construction of the bin-based genetic map. (C) Original called SNP data from GATK program, without filtering and binning. ** 0: SNP from R500; 2: SNP from L58; 1: Heterozygous SNP; −1:Missing value.

**Supplemental Table S8.** (A) Amino acid identity among *A. thaliana* and *B. rapa* R500 *PAP* genes. (B) Amino acid polymorphisms in the coding regions of the R500 and L58 alleles of *PAP2(A03)*.

**Supplemental File S1.** LTR Libraries with chromosomal location noted. File is in Fasta format.

**Supplemental File S2.** R Code used for seed coat color analysis.

## Literature Cited

Balasubramanian, S., Sureshkumar, S., Lempe, J., and Weigel, D. (2006). Potent induction of *Arabidopsis thaliana* flowering by elevated growth temperature. PLoS Genet. 2: e106. doi:10.1371/journal.pgen.0020106

Balasubramanian, S., Schwartz, C., Singh, A., Warthmann, N., Kim, M.C., Maloof, J.N., Loudet, O., Trainer, G.T., Dabi, T., Borevitz, J.O., Chory, J., and Weigel, D. (2009). QTL mapping in new *Arabidopsis thaliana* Advanced Intercross-Recombinant Inbred Lines. PLoS ONE 4: e4318.

Belser, C., Istace, B., Denis, E., Dubarry, M., Baurens, F.-C., Falentin, C., Genete, M., Berrabah, W., Chèvre, A.-M., Delourme, R., Deniot, G., Denoeud, F., Duffé, P., Engelen, S., Lemainque, A., Manzanares-Dauleux, M., Martin, G., Morice, J., Noel, B., Vekemans, X., D’Hont, A., Rousseau-Gueutin, M., Barbe, V., Cruaud, C., Wincker, P., and Aury, J.-M. (2018). Chromosome-scale assemblies of plant genomes using nanopore long reads and optical maps. Nat. Plants 4: 879–887. doi:10.1038/s41477-018-0289-4

Bolger, A.M., Lohse, M., and Usadel, B. (2014). Trimmomatic: a flexible trimmer for Illumina sequence data. Bioinformatics 30: 2114–2120. doi:10.1093/bioinformatics/btu170

Borevitz, J.O., Xia, Y., Blount, J., Dixon, R.A., and Lamb, C. (2000). Activation tagging identifies a conserved MYB regulator of phenylpropanoid biosynthesis. Plant Cell 12: 2383–2393. doi:10.1105/tpc.12.12.2383

Broman, K.W., Wu, H., Sen, S., and Churchill, G.A. (2003). R/qtl: QTL mapping in experimental crosses. Bioinformatics 19: 889–890.

Campbell, M.S., Law, M., Holt, C., Stein, J.C., Moghe, G.D., Hufnagel, D.E., Lei, J., Achawanantakun, R., Jiao, D., Lawrence, C.J., Ware, D., Shiu, S.-H., Childs, K.L., Sun, Y., Jiang, N., and Yandell, M. (2014). MAKER-P: A tool kit for the rapid creation, management, and quality control of plant genome annotations. Plant Physiol. 164: 513–524. doi:10.1104/pp.113.230144

Cominelli, E., Gusmaroli, G., Allegra, D., Galbiati, M., Wade, H.K., Jenkins, G.I., and Tonelli, C. (2008). Expression analysis of anthocyanin regulatory genes in response to different light qualities in *Arabidopsis thaliana*. J. Plant Physiol. 165: 886–894. doi:10.1016/j.jplph.2007.06.010

Danecek, P., Auton, A., Abecasis, G., Albers, C.A., Banks, E., DePristo, M.A., Handsaker, R.E., Lunter, G., Marth, G.T., Sherry, S.T., McVean, G., Durbin, R., and Group, G.P.A. (2011). The variant call format and VCFtools. Bioinformatics 27: 2156–2158. doi:10.1093/bioinformatics/btr330

Ellinghaus, D., Kurtz, S., and Willhoeft, U. (2008). LTRharvest, an efficient and flexible software for de novo detection of LTR retrotransposons. BMC Bioinformatics 9: 18. doi:10.1186/1471-2105-9-18

Gonda, I., Ashrafi, H., Lyon, D.A., Strickler, S.R., Hulse-Kemp, A.M., Ma, Q., Sun, H., Stoffel, K., Powell, A.F., Futrell, S., Thannhauser, T.W., Fei, Z., Van Deynze, A.E., Mueller, L.A., Giovannoni, J.J., and Foolad, M.R. (2019). Sequencing-based bin map construction of a tomato mapping population, facilitating high-resolution quantitative trait loci detection. Plant Genome 12: 180010. doi:10.3835/plantgenome2018.02.0010

Haas, B.J., Papanicolaou, A., Yassour, M., Grabherr, M., Blood, P.D., Bowden, J., Couger, M.B., Eccles, D., Li, B., Lieber, M., MacManes, M.D., Ott, M., Orvis, J., Pochet, N., Strozzi, F., Weeks, N., Westerman, R., William, T., Dewey, C.N., Henschel, R., LeDuc, R.D., Friedman, N., and Regev, A. (2013). De novo transcript sequence reconstruction from RNA-seq using the Trinity platform for reference generation and analysis. Nat. Protoc. 8: 1494–1512. doi:10.1038/nprot.2013.084

Han, Y., and Wessler, S.R. (2010). MITE-Hunter: a program for discovering miniature inverted-repeat transposable elements from genomic sequences. Nuc. Acids Res. 38. doi:10.1093/nar/gkq862

Hohmann, N., Wolf, E.M., Lysak, M.A., and Koch, M.A. (2015). A time-calibrated road map of Brassicaceae species radiation and evolutionary history. Plant Cell 27: 2770–2784. doi:10.1105/tpc.15.00482

Iniguez-Luy, F.L., Lukens, L., Farnham, M.W., Amasino, R.M., and Osborn, T.C. (2009). Development of public immortal mapping populations, molecular markers and linkage maps for rapid cycling *Brassica rapa* and *B. oleracea*. Theor. Appl. Genet. 119: 31–43.

Jurka, J., Kapitonov, V.V., Pavlicek, A., Klonowski, P., Kohany, O., and Walichiewicz, J. (2005). Repbase Update, a database of eukaryotic repetitive elements. Cytogenet. Genome Res. 110: 462–467.

Kebede, B., Cheema, K., Greenshields, D.L., Li, C., Selvaraj, G., and Rahman, H. (2012). Construction of genetic linkage map and mapping of QTL for seed color in *Brassica rapa*. Genome 55: 13–823. doi:10.1139/g2012-066

Korf, I. (2004). Gene finding in novel genomes. BMC Bioinformatics 5: 59. doi:10.1186/1471-2105-5-59

Langmead, B., and Salzberg, S.L. (2012). Fast gapped-read alignment with Bowtie 2. Nat. Meth. 9: 357–359. doi:10.1038/nmeth.1923

Li, H., and Durbin, R. (2009). Fast and accurate short read alignment with Burrows–Wheeler transform. Bioinformatics 25: 1754–1760. doi:10.1093/bioinformatics/btp324

Li, N., Wu, H., Ding, Q., Li, H., Li, Z., Ding, J., and Li, Y. (2018). The heterologous expression of Arabidopsis PAP2 induces anthocyanin accumulation and inhibits plant growth in tomato. Funct. Integr. Genomics 18: 341–353. doi:10.1007/s10142-018-0590-3

Li, X., Chen, L.-Q., Hong, M., Zhang, Y., Zu, F., Wen, J., Yi, B., Ma, C., Shen, J., Tu, J., and Fu, T. (2012). A large insertion in bHLH transcription factor *BrTT8* resulting in yellow seed coat in *Brassica rapa*. PLoS ONE 7: e44145. doi:10.1371/journal.pone.0044145

Lincoln, S.E., and Lander, E.S. (1992). Systematic detection of errors in genetic linkage data. Genomics 14: 604–610.

Margarido, G.R.A., Souza, A.P., and Garcia, A.A.F. (2007). OneMap: software for genetic mapping in outcrossing species. Hereditas 144: 78–79. doi:10.1111/j.2007.0018-0661.02000.x

Markelz, R.J.C., Covington, M.F., Brock, M.T., Devisetty, U.K., Kliebenstein, D.J., Weinig, C., and Maloof, J.N. (2017). Using RNA-seq for genomic scaffold placement, correcting assemblies, and genetic map creation in a common *Brassica rapa* mapping population. G3 7: 2259–2270. doi:10.1534/g3.117.043000

McKenna, A., Hanna, M., Banks, E., Sivachenko, A., Cibulskis, K., Kernytsky, A., Garimella, K., Altshuler, D., Gabriel, S., Daly, M., and DePristo, M.A. (2010). The Genome Analysis Toolkit: A MapReduce framework for analyzing next-generation DNA sequencing data. Genome Res. 20: 1297–1303. doi:10.1101/gr.107524.110

Mitchell, A.L.e., Attwood, T.K., Babbitt, P.C., Blum, M., Bork, P., Bridge, A., Brown, S.D., Chang, H.Y., El-Gebali, S., Fraser, M.I., Gough, J., Haft, D.R., Huang, H., Letunic, I., Lopez, R., Luciani, A., Madeira, F., Marchler-Bauer, A., Mi, H., Natale, D.A., Necci, M., Nuka, G., Orengo, C., Pandurangan, A.P., Paysan-Lafosse, T., Pesseat, S., Potter, S.C., Qureshi, M.A., Rawlings, N.D., Redaschi, N., Richardson, L.J., Rivoire, C., Salazar, G.A., Sangrador-Vegas, A., Sigrist, C.J.A., Sillitoe, I., Sutton, G.G., Thanki, N., Thomas, P.D., Tosatto, S.C.E., Yong, S.Y., and Finn, R.D. (2019). InterPro in 2019: improving coverage, classification and access to protein sequence annotations. Nucleic Acids Res. 47: D351–D360. doi:10.1093/nar/gky1100

Moriya, Y., Itoh, M., Okuda, S., Yoshizawa, A.C., and Kanehisa, M. (2007). KAAS: an automatic genome annotation and pathway reconstruction server. Nucleic Acids Res. 35: W182–185. doi:10.1093/nar/gkm321

Nesi, N., Debeaujon, I., Jond, C., Pelletier, G., Caboche, M., and Leiniec, L. (2000). The *TT8* gene encodes a basic helix-loop-helix domain protein required for expression of *DFR* and *BAN* genes in Arabidopsis siliques. Plant Cell 12: 1863–1878.

Ou, S., and Jiang, N. (2018). LTR_retriever: A highly accurate and sensitive program for identification of Long Terminal Repeat retrotransposons. Plant Physiol. 176: 1410–1422. doi:10.1104/pp.17.01310

Ouellette, L.A., Reid, R.W., Blanchard, S.G., and Brouwer, C.R. (2017). LinkageMapView—rendering high-resolution linkage and QTL maps. Bioinformatics 34: 306–307. doi:10.1093/bioinformatics/btx576

Padmaja, L.K., Agarwal, P., Gupta, V., Mukhopadhyay, A., Sodhi, Y.S., Pental, D., and Pradhan, A.K. (2014). Natural mutations in two homoeologous *TT8* genes controll yellow seed coat trait in allotetraploid *Brassica juncea* (AABB). Theor. Appl. Genet. 127: 339–347.

Parkin, I.A.P., Gulden, S.M., Sharpe, A.G., Lukens, L., Trick, M., Osborn, T.C., and Lydiate, D.J. (2005). Segmental structure of the *Brassica napus* genome based on comparative analysis with *Arabidopsis thaliana*. Genetics 171: 765–781.

Pertea, M., Pertea, G.M., Antonescu, C.M., Chang, T.-C., Mendell, J.T., and Salzberg, S.L. (2015). StringTie enables improved reconstruction of a transcriptome from RNA-seq reads. Nat. Biotechnol. 33: 290–295. doi:10.1038/nbt.3122

Qi, X., An, H., Ragsdale, A.P., Hall, T.E., Gutenkunst, R.N., Pires, J.C., and Barker, M.S. (2017). Genomic inferences of domestication events are corroborated by written records in *Brassica rapa*. Mol. Ecol. 26: 3373–3388. doi:10.1111/mec.14131

Qi, X., An, H., Hall, T.E., Di, C., Blischak, P.D., Pires, J.C., and Barker, M.S. (2019). Genes derived from ancient polyploidy have higher genetic diversity and are associated with domestication in *Brassica rapa*. bioRxiv. doi:https://doi.org/10.1101/842351

R Core Team (2018). R: A language and environment for statistical computing. Vienna, Austria. URL https://www.R-project.org/.

Rahman, M., Mamidi, S., and McClean, P. (2014). Quantitative trait loci mapping of seed colour, hairy leaf, seedling anthocyanin, leaf chlorosis and days to flowering in F2 population of *Brassica rapa* L. Plant Breeding 133: 381–389.

Schranz, M., Lysak, M., and Mitchell-Olds, T. (2006). The ABC’s of comparative genomics in the Brassicaceae: building blocks of crucifer genomes. Trends Plant Sci. 11: 535–542.

Simão, F.A., Waterhouse, R.M., Ioannidis, P., Kriventseva, E.V., and Zdobnov, E.M. (2015). BUSCO: assessing genome assembly and annotation completeness with single-copy orthologs. Bioinformatics 31: 3210–3212.

Slabaugh, E., Desai, J.S., Sartor, R.C., Lawas, L.M.F., Krishna Jagadish, S.V., and Doherty, C.J. (2019). Analysis of differential gene expression and alternative splicing is significantly influenced by choice of reference genome. RNA 25: 669–684. doi:10.1261/rna.070227.118

Smit, A.F.A., and Hubley, R. (2008). RepeatModeler Open-1.0. http://www.repeatmasker.org.

Smit, A.F.A., Hubley, R., and Green, P. (1996). RepeatMasker Open-3.0. http://www.repeatmasker.org.

Stanke, M., and Waack, S. (2003). Gene prediction with a hidden Markov model and a new intron submodel. Bioinformatics 19 Suppl 2: ii215–225. doi:10.1093/bioinformatics/btg1080

Tang, H., Woodhouse, M.R., Cheng, F., Schnable, J.C., Pedersen, B.S., Conant, G., Wang, X., Freeling, M., and Pires, J.C. (2012). Altered patterns of fractionation and exon deletions in *Brassica rapa* support a two-step model of paleohexaploidy. Genetics 190: 1563–1574. doi:10.1534/genetics.111.137349

Town, C.D., Cheung, F., Maiti, R., Crabtree, J., Haas, B.J., Wortman, J.R., Hine, E.E., Althoff, R., Arbogast, T.S., Tallon, L.J., Vigouroux, M., Trick, M., and Bancroft, I. (2006). Comparative genomics of *Brassica oleracea* and *Arabidopsis thaliana* reveal gene loss, fragmentation, and dispersal after polyploidy. Plant Cell 18: 1348–1359.

Wang, X., Wang, H., Wang, J., Sun, R., Wu, J., Liu, S., Bai, Y., Mun, J.-H., Bancroft, I., Cheng, F., Huang, S., Li, X., Hua, W., Wang, J., Wang, X., Freeling, M., Pires, J.C., Paterson, A.H., Chalhoub, B., Wang, B., Hayward, A., Sharpe, A.G., Park, B.-S., Weisshaar, B., Liu, B., Li, B., Liu, B., Tong, C., Song, C., Duran, C., Peng, C., Geng, C., Koh, C., Lin, C., Edwards, D., Mu, D., Shen, D., Soumpourou, E., Li, F., Fraser, F., Conant, G., Lassalle, G., King, G.J., Bonnema, G., Tang, H., Wang, H., Belcram, H., Zhou, H., Hirakawa, H., Abe, H., Guo, H., Wang, H., Jin, H., Parkin, I.A.P., Batley, J., Kim, J.-S., Just, J., Li, J., Xu, J., Deng, J., Kim, J.A., Li, J., Yu, J., Meng, J., Wang, J., Min, J., Poulain, J., Wang, J., Hatakeyama, K., Wu, K., Wang, L., Fang, L., Trick, M., Links, M.G., Zhao, M., Jin, M., Ramchiary, N., Drou, N., Berkman, P.J., Cai, Q., Huang, Q., Li, R., Tabata, S., Cheng, S., Zhang, S., Zhang, S., Huang, S., Sato, S., Sun, S., Kwon, S.-J., Choi, S.-R., Lee, T.-H., Fan, W., Zhao, X., Tan, X., Xu, X., Wang, Y., Qiu, Y., Yin, Y., Li, Y., Du, Y., Liao, Y., Lim, Y., Narusaka, Y., Wang, Y., Wang, Z., Li, Z., Wang, Z., Xiong, Z., and Zhang, Z. (2011). The genome of the mesopolyploid crop species *Brassica rapa*. Nat. Genet. 43: 1035–1039. doi:10.1038/ng.919

Wang, Y., Xiao, L., Guo, S., An, F., and Du, D. (2016). Fine mapping and whole-genome resequencing identify the seed coat color gene in *Brassica rapa*. PLoS ONE 11: e0166464. doi:10.1371/journal.pone.0166464

Wick, R.R., Schultz, M.B., Zobel, J., and Holt, K.E. (2015). Bandage: interactive visualization of de novo genome assemblies. Bioinformatics 31: 3350–3352. doi:10.1093/bioinformatics/btv383

Xu, Z., and Wang, H. (2007). LTR_FINDER: an efficient tool for the prediction of full-length LTR retrotransposons. Nucleic Acids Res. 35: W265–268. doi:10.1093/nar/gkm286

Zhang, L., Cai, X., Wu, J., Liu, M., Grob, S., Cheng, F., Liang, J., Cai, C., Liu, Z., Liu, B., Wang, F., Li, S., Liu, F., Li, X., Cheng, L., Yang, W., Li, M.-h., Grossniklaus, U., Zheng, H., and Wang, X. (2018). Improved *Brassica rapa* reference genome by single-molecule sequencing and chromosome conformation capture technologies. Hort. Res. 5: 50. doi:10.1038/s41438-018-0071-9

Zhang, X., Zhang, K., Wu, J., Guo, N., Liang, J., Wang, X., and Cheng, F. (2020). QTL-Seq and sequence assembly rapidly mapped the gene *BrMYBL2.1* for the purple trait in *Brassica rapa*. Sci. Rep. 10: 2328. doi:10.1038/s41598-020-58916-5

Zhao, H., Basu, U., Kebede, B., Qu, C., Li, J., and Rahman, H. (2019). Fine mapping of the major QTL for seed coat color in *Brassica rapa* var. Yellow Sarson by use of NIL populations and transcriptome sequencing for identification of the candidate genes. PLoS ONE 14: e0209982. doi:10.1371/journal.pone.0209982

Zhao, J., Artemyeva, A., Del Carpio, D.P., Basnet, R.K., Zhang, N., Gao, J., Li, F., Bucher, J., Wang, X., Visser, R.G.F., and Bonnema, G. (2010). Design of a *Brassica rapa* core collection for association mapping studies. Genome 53: 884–898.

