## Supplemental Figure S1 for "Genetic and Genomic Resources to Study Natural Variation in *Brassica rapa*"

**Supplemental File S1.** LTR Libraries with chromosomal location noted. File is in Fasta format.

**Supplemental File S2.** R Code used for seed coat color analysis.

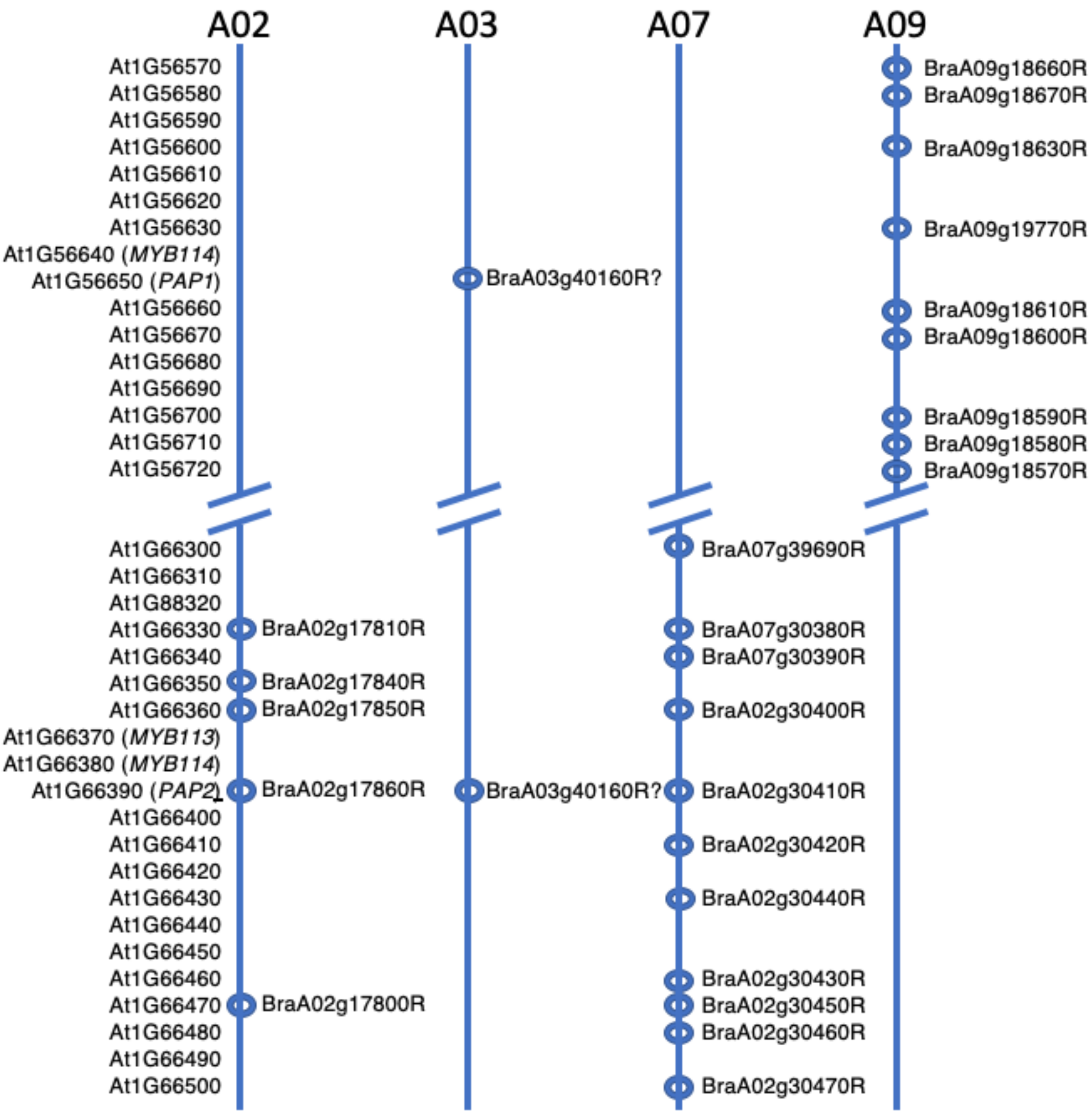


**Supplemental Figure S1A.** Microsynteny analysis of members of the *PAP* Gene Family in Arabidopsis and *Brassica rapa*.

*B. rapa* genes retained after fractionation in each of the genomic regions syntenic to the Arabidopsis PAP genes are indicated.

```
Arabidopsis_PAP1          MEGSSKGLRKGAWTTEEDSLLRQCINKYGEGKWHQVPVRAGLNRCRKSCRLRWLNYLKPS  60
Arabidopsis_PAP2          MEGSSKGLRKGAWTAEEDSLLRLCIDKYGEGKWHQVPLRAGLNRCRKSCRLRWLNYLKPS  60
BraA02g17860R_L58_A02     MEGSPKGLRKGAWTAEEDSLLRQCIDKYGEGKWHQVPLRAGLNRCRKSCRLRWLNYLKPS  60
BraA02g17860R_R500_A02    MEGSPKGLRKGAWTAEEDSLLRQCIDKYGEGKWHQVPLRAGLNRCRKSCRLRWLNYLKPS  60
BraA07g30410R_L58_A07     MEGSSQGLKKGAWTAEEDNLLRQCIDKYGEGKWHQVPLRAGLNRCRKSCRLRWLNYLKPS  60
BraA07g30410R_R500_A07    MEGSSQGLKKGAWTAEEDNLLRQCIDKYGEGKWHQVPLRAGLNRCRKSCRLRWLNYLKPS  60
BraA03g40160R_A03_L58     MEDSSKGLTKGAWTAEEDSLLRRCIDKYGEGKWHQVPLRAGLNRCRKSCRLRWLNYLKPN  60
BraA03g40160R_A03_R500    MEDSSKGLTKGAWTAEEDSLLRRCIDKYGEGKWHQIPLRAGLNRCRKSCRLRWLNYLKPT  60
                          **.* :** *****:***.*** **:*********:*:*********************.

Arabidopsis_PAP1          IKRGKLSSDEVDLLLRLHRLLGNRWSLIAGRLPGRTANDVKNYWNTHLSKKHE-PCCKIK  119
Arabidopsis_PAP2          IKRGRLSNDEVDLLLRLHKLLGNRWSLIAGRLPGRTANDVKNYWNTHLSKKHESSCCKSK  120
BraA02g17860R_L58_A02     IKKGKLSSDEVDLLLRLHKLLGNRWSLIAGRLPGRTANDVKNYWNTHLSKKHE-PGCNTK  119
BraA02g17860R_R500_A02    IKKGKLSSDEVDLLLRLHKLLGNRWSLIAGRLPGRTANDVKNYWNTHLSKKHE-PGCNTK  119
BraA07g30410R_L58_A07     IKRGKLNSDEVDLLIRLHKLLGNRWSLIAGRLPGRTANDVKNYWNTHLSKKHE-PGCKTQ  119
BraA07g30410R_R500_A07    IKRGKLNSDEVDLLIRLHKLLGNRWSLIAGRLPGRTANDVKNYWNTHLSKKHE-PGCKTQ  119
BraA03g40160R_A03_L58     IKRGKLSSDEVDLLLRLHKLLGNRWSLIAGRLPGRTANDIKNYWNTHLSKKHE-PCCKTK  119
BraA03g40160R_A03_R500    IKRGKLSSDEVDLLLRLHKLLGNRWSLIAGRLPGRTANDIKNYWNTHLSKKHE-PCCKTK  119
                          **:*:*..******:***:*******************:*************   *: :

Arabidopsis_PAP1          MKKRDITPIPTTPALKNNVYKPRPRSFTVNNDCNHLNAPPKVDVNPPCLGLN-INNVCDN  178
Arabidopsis_PAP2          MKKKNIISPPTTPVQKIGVFKPRPRSFSVNNGCSHLNGLPEVDLIPSCLGLK-KNNVCEN  179
BraA02g17860R_L58_A02     MRKRNIPCSSTQPAQKNEVLKPRPRSFTVNNGCSHFNGKPKVDVIPLFLGVNNTNNVCEN  179
BraA02g17860R_R500_A02    MRKRNIPCSSTQPAQKNEVLKPRPRSFTVNNGCSHFNGKPKVDVIPLFLGVNNTNNVCEN  179
BraA07g30410R_L58_A07     MKKRNIPCSYTTPAQKIDVFKPRPRSFTVNSGCSHNNGMPEAGIVPLCLGHNDTNNVSEN  179
BraA07g30410R_R500_A07    MKKRNIPCSYTTPAQKIDVFKPRPRSFTVNSGCSHNNGMPEAGIVPLCLGHNDTNNVSEN  179
BraA03g40160R_A03_L58     MKKRNVTFSSSTTPAQKIDVFKPRPRLFTVNNGCSHLHGLPEVDVVPPCLGLNNINNVCEN  179
BraA03g40160R_A03_R500    MKKRNVTFSSSTTPAQKIDVFKPRPRLFTVNDGCSHLHGLPEVDVVPPCVGLNNINNVCEN  179
                          *:*:::    * *. *  * ***** *:**..*.* :. *:..: *  :* :  ***.:*

Arabidopsis_PAP1          SIIYNKDKKKDQLVNNLIDGDNMWLEKFLEESQEVDILVPEATTTEKGDTLAFDVDQLWS  238
Arabidopsis_PAP2          SITCNKDDEKDDFVNNLMNGDNMWLENLLGENQEADAIVPEATTAEHGATLAFDVEQLWS  239
BraA02g17860R_L58_A02     SITYKKDAEKYELVNNLMDGENMWWKSLLEESQEPDAIVPESTETEKLATSAFDVEQLWN  239
BraA02g17860R_R500_A02    SITYKKDAEKYELVNNLMDGENMWWKSLLEESQEPDAIVPESTETEKLATSAFDVEQLWN  239
BraA07g30410R_L58_A07     IITCNKDDDKSELVSHLMDGQNRWWESLLDESQDPAALFPETTAIKKGATSAFDVEQLWS  239
BraA07g30410R_R500_A07    IITCNKDDDKSELVSHLMDGQNRWWESLLDESQDPAALFPETTAI---------------  224
BraA03g40160R_A03_L58     SMTCNKAGEKYELYSNLMDGENMWWESLLEESKQPDGLVPKGTATKKGATFAFDVEQLWN  239
BraA03g40160R_A03_R500    SMTCNKAGEKYELFSNLMDGENMWWESLLEQSKQPDGLVPKGTATKKGATFAFDVEQLWN  239
                           :  :*  .* :: .:*::*:* * :.:* :.::   :.*: *

Arabidopsis_PAP1          LFDGETVKFD       248
Arabidopsis_PAP2          LFDGETVELD       249
BraA02g17860R_L58_A02     LLDGETVELD       249
BraA02g17860R_R500_A02    LLDGETVELD       249
BraA07g30410R_L58_A07     LLDGETGT--       247
BraA07g30410R_R500_A07    ----------       224
BraA03g40160R_A03_L58     MLDGETVELD       249
BraA03g40160R_A03_R500    MLDGETVELD       249
```

**Supplemental Figure S1B.** Multiple alignment of PAP proteins of *Arabidopsis* and *Brassica rapa.* Amino acid differences between the R500 and L58 PAP(A03) variants are colored, with highly conserved residues in red and dissimilar residues in yellow or cyan. *, :, and . indicates fully, strongly, and weakly conserved, respectively.

```
R500 1    ATGGAGGATTCGTCCAAAGGGTTGACCAAAGGTGCATGGACGGCTGAAGAAGACAGTCTCTTGAGGCGATGCATTGATAAGTATGGAGAAGGCAAATGGC 100
L58  1    ATGGAGGATTCGTCCAAAGGGTTGACAAAAGGTGCATGGACCGCTGAAGAAGACAGTCTCTTGAGGCGATGCATTGATAAGTATGGAGAAGGCAAATGGC 100

     101  ATCAAATTCCTTTAAGAGCTGGTATGTTACTTTTTTTTCTTT-TGT-ACACACACAT-------------ATCTGTATACATATAATTAATCACTACGAA 185
     101  ATCAAGTTCCTTTAAGAGCTGGTATGTTACTTTTTTT-CTT-CTG-CACACACACATATATGTACATATGATCTGTATACATATAATTAATCACTATGAA 197

     186  AAAC-TTCTTTC-TCTTTGTCTTCTACAGTACTACTCGGA-GAATTAATTAACACATGGCTGCACAAAACAAAGTTTTTCTTTTTGTCATAATGAACAAA 282
     198  AAA-TTTCTTT-ATCTTTG---TCTAC--TA-T--TCGG-GGAATTAATTAACACATGGCTGCACAAAGCAAAGTTTTTCTTTTTGTCATAATGAACAAA 286

     283  TCTTTGATTCATGCTTTATGCGGTTGATCATGAAAAAA-CATTCATGTTGTTCATATTAATTATGTCGCACTTAAACGAAGATCTATAATTAAGACTTTT 381
     287  TCTTTGACTCATGCTTTATGCGGTTGTTCATGAAAAAAACATTCATGTTGTTCATATTAATTATGTCGCACTTAAACGAAGATCTATAATTAAGGCTTTT 386

     382  ACTTTCA-CTGACAAAGCGAAAAGATACCAATAATTTTTTTG-GAC-TGTCCTTTAGTACATGAATTCAATGACATTTTCTGTACGAGCACGTGTCTTTG 478
     387  ACTTTC-GCTGACAAAGCGAAAAGATACCAACAATTTTTTT-CGA-TTGTCCTTTAGTACATGAATTCAGTGACATTTTCTGTACGAACACGTGTCTTTG 483

     479  TGTGGCAATAATT-ATATATAATTTCTGTTAGTGTGATTCTTCGTGATAAAATTATTGGTTTTGTAGGGCTTAATAGGTGTAGGAAGAGTTGTAGACTAA 577
     484  TGTGG-A--AAT-AATATATAATTTCTGTTAGTGTTATTCTTCGTGATAAAATTATTGGTTTTGTAGGGCTTAATAGGTGTAGGAAGAGTTGTAGACTAA 579

     578  GATGGCTGAACTATTTGAAGCCAACTATCAAGAGAGGAAAACTTAGCTCTGATGAAGTTGATCTTCTTCTCCGTCTTCATAAGCTTTTAGGAAACAGGTT 677
     580  GATGGCTGAACTATTTGAAGCCAAATATCAAGAGAGGAAAACTTAGCTCTGATGAAGTTGATCTTCTTCTCCGTCTTCATAAGCTTTTAGGAAACAGGTT 679

     678  TGTATTCTTAAGACAAAAATTCAACTTTGTT-TCTTGTCTAATGATCCATAAGA--------------TATATATATGTATATCCAAATCGTTCAAATGC 762
     680  TGTATTCTTAAGACAAAAATTCAACTT-GT-ATCTTGTCTAATGATCCATAAGATATATATATATATATATATATATGTATATCCAAATCGTTCAAATGC 777

     763  ATGCTTAGGTGGTCTTTAATTGCTGGTAGACTACCCGGTCGGACCGCTAATGATATCAAGAATTACTGGAACACCCATCTGAGCAAGAAACATGAACCAT 862
     778  ATGCTTAGGTGGTCTTTAATTGCTGGTAGACTACCCGGTCGGACCGCTAATGATATCAAGAATTACTGGAACACCCATCTGAGCAAGAAACATGAACCAT 877

     863  GTTGTAAGACCAAGATGAAGAAGAGAAACGTTACATTCTCTTCTACCACACCCGCCCAAAAAATCGACGTTTTCAAACCTCGACCTCGACTCTTCACCGT 962
     878  GTTGTAAGACCAAGATGAAGAAGAGAAACGTTACATTCTCTTCTACCACACCCGCCCAAAAAATCGACGTTTTCAAACCTCGACCTCGACTCTTCACCGT 977

     963  TAACGATGGCTGCAGCCATCTCCATGGCCTGCCAGAAGTTGACGTTGTTCCTCCATGCGTTGGACTCAACAACATTAATAATGTCTGTGAAAATAGTATG 1062
     978  TAACAATGGCTGCAGCCATCTCCATGGCCTGCCAGAAGTTGACGTTGTTCCTCCATGCCTTGGACTCAACAACATTAATAATGTCTGTGAAAATAGTATG 1077

     1063 ACATGTAACAAAGCTGGGGAGAAGTATGAACTTTTTAGTAATTTAATGGATGGAGAGAATATGTGGTGGGAGAGTTTGCTAGAGCAGAGCAAACAGCCTG 1162
     1078 ACATGTAACAAAGCTGGGGAGAAGTATGAACTTTATAGTAATTTAATGGATGGAGAGAATATGTGGTGGGAGAGTTTGCTAGAGGAGAGCAAACAGCCTG 1177

     1163 ACGGGCTCGTTCCAAAAGGTACGGCAACAAAAAAGGGGGCAACCTTTGCGTTTGACGTTGAGCAACTTTGGAATATGTTGGATGGAGAGACTGTAGAACT 1262
     1178 ACGGGCTCGTTCCAAAAGGTACGGCAACAAAAAAGGGGGCAACCTTTGCGTTTGACGTTGAGCAACTTTGGAATATGTTGGATGGAGAGACTGTAGAACT 1277

     1263 TGATTAG 1269
     1278 TGATTAG 1284
```

**Supplemental Figure S1C.** Nucleotide sequence alignment of R500 (upper) and L58 (lower) alleles of the *PAP2(A03)* (BraA03g40160R) gene. Exonic sequences are highlighted in cyan and intronic sequences are not highlighted. Nucleotide polymorphisms are indicated by red, with substitutions indicated by # and indels indicated by filled boxes. Asterisks mark 10 bp intervals.
