## Supplemental File S2 for "Genetic and Genomic Resources to Study Natural Variation in *Brassica rapa*"

**Supplemental File S2. R Code used for seed coat color analysis.**

install.packages("qtl");

library(qtl)

#read the csv files

L58_SCC <-read.cross("csv","your path to csv file","File name")

#convert to RIL population

L58_SCC <- convert2riself(L58_SCC)

\# summary/plot here

summary(L58_SCC)

plot(L58_SCC)

# remove the markers with duplicate location before analysis

L58_SCC <- jittermap(L58_SCC)

\# calculate the geno prob

L58_SCC <- calc.genoprob(L58_SCC,step=0.5,error.prob = 0.001)

\# by default the trait is in column 1

L58_SCC.em <-scanone(L58_SCC, pheno.col=2, method="em")

\# plot and summary

plot (L58_SCC.em, ylab="LOD")

summary (L58_SCC.em)

#LOD threshold

operm <- scanone(L58_SCC, n.perm=1000, verbose=FALSE);

summary(operm, alpha=c(0.05, 0.01));

\# for CIM mapping, number of co-factors depends on scanone output, for example, how many QTL identified. Windows size of 20 was used in current analysis

L58_SCC.cim <-cim(L58_SCC,pheno.col=2,n.marcovar=2, window=20)

\# summary/plot

plot(L58_SCC.cim)

summary(L58_SCC.cim)

\# statistically significant interval was determined by using lodint at 1.5.

lodint(L58_SCC.cim,A09,125,expandtomarkers = TRUE)

lodint(L58_SCC.cim,A03,195,expandtomarkers = TRUE)
